# Distinct classes of lagging chromosome underpin age-related oocyte aneuploidy in mouse

**DOI:** 10.1101/2021.01.30.428896

**Authors:** Aleksandar I. Mihajlović, Jenna Haverfield, Greg FitzHarris

## Abstract

Chromosome segregation errors that cause oocyte aneuploidy increase in frequency with maternal age and are considered a major contributing factor of age-related fertility decline in females. A common age-associated chromosome segregation phenomenon in oocytes is the lagging anaphase chromosome, but whether anaphase laggards actually missegregate and cause aneuploidy is unclear. Here we show unexpectedly that lagging chromosomes in mouse oocytes comprise two mechanistically distinct classes of motion that we refer to as ‘Class-I’ and ‘Class-II’. We use imaging approaches and mechanistic interventions to dissociate the two classes, and find that whereas Class-II laggards are benign, Class-I laggards can directly cause aneuploidy. Most notably, a controlled prolongation of meiosis-I specifically lessens Class-I lagging to prevent aneuploidy. Our data thus reveal lagging chromosomes to be a cause of age-related aneuploidy in mouse oocytes and suggest that manipulating the cell cycle could increase the yield of useful oocytes in some contexts.

## INTRODUCTION

Failure to correctly segregate chromosomes during cell division leads to aberrations in their number, termed aneuploidy. In somatic cells, aneuploidy causes tumorigenesis and is considered a hallmark of cancer (Hanahan and Weinberg, 2011). In the germ line, it leads to developmental breakdown and reproductive failure (Hassold and Hunt, 2001; Ioannou et al., 2019; Nagaoka et al., 2012; Templado et al., 2013). Specifically, chromosome segregation errors increase dramatically with maternal age during meiosis-I (MI) in mammals, including humans and mice (Greaney et al., 2018; Mihajlović and FitzHarris, 2018; Webster and Schuh, 2017), producing aneuploid Metaphase-II (Met-II) eggs that are developmentally compromised. Despite over a decade of intense study, the cellular explanation for the age-related increase in oocyte aneuploidy remains incomplete.

Proper chromosome segregation in MI involves separation of homologous chromosomes that exist as pairs of sister chromatids in a structure called a bivalent (Herbert et al., 2015), to either the egg or the polar body. A well-established age-related lesion leading to chromosome segregation error is a progressive loss of chromosome cohesion (Chiang et al., 2010, 2011; Duncan et al., 2012; Lister et al., 2010; Merriman et al., 2012) that allows premature separation of bivalents into pairs of sister chromatids called univalents (Sakakibara et al., 2015; Zielinska et al., 2015). Although this is irrefutably a major contributor to age-related oocyte aneuploidy, the reported frequency of segregation error caused by univalent formation is insufficient to fully account for all observed oocyte aneuploidy (Chiang et al., 2010; Sakakibara et al., 2015; Yun et al., 2014; Zielinska et al., 2015). Thus, the causes of age-related oocyte aneuploidy remain to be fully resolved.

One well-known phenomenon that could contribute to aneuploidy is the so-called lagging anaphase chromosome, wherein a chromosome ‘lags’ behind the others headed towards the same spindle pole (Ganem and Pellman, 2012; Salmon et al., 2005). Well-studied in somatic cells, laggards are a result of aberrant ‘merotelic’ attachment of kinetochores to both spindle poles simultaneously (Cimini et al., 2001, 2002). Whilst it is intuitive that this puts them at risk of missegregation, it remains unclear how often they actually cause aneuploidy (Cimini et al., 2001; Thompson and Compton, 2011). In oocytes, a substantial age-related increase in the incidence of laggards in anaphase of MI has been reported, alluding to their involvement in aneuploidy (Chiang et al., 2010; Lister et al., 2010; Nakagawa and FitzHarris, 2017; Yun et al., 2014). However, their reported frequency tends to far exceed the rate of age-related aneuploidy and so the consequence of lagging is yet unclear.

Here we set out to uncover the role of the lagging chromosome in oocyte aneuploidy. We report two mechanistically distinct and experimentally separable classes of lagging chromosome motions, with differing implications for the fidelity of chromosome segregation. Our experiments show that whereas one class is largely benign, the other directly contributes to aneuploidy.

## RESULTS

### Two classes of lagging chromosomes in oocytes

To investigate the relationship between anaphase lagging and aneuploidy we first assessed each in oocytes from CD1 mice at 2-3 months and 16 months of age (hereinafter referred to as ‘young’ and ‘aged’ oocytes). *In situ* chromosome counting (Duncan et al., 2009), revealed an increase in the aneuploidy level from 1.7% in young to 23.1% in aged oocytes (Fig. S1; P<0.001), similar to other studies (Chiang et al., 2010; Duncan et al., 2009; Pan et al., 2008; Shomper et al., 2014). To assess the incidence of anaphase lagging during MI, chromosomes and centromeres of young and aged germinal vesicle (GV)-stage oocytes were labelled with fluorescently tagged histone 2B (H2B-RFP) and major satellite probe (Maj.Sat.-mClover) respectively, and live-imaged until Met-II (Fig. 1a). The incidence of lagging chromosomes increased from 21.7% in young to 66.7% in aged oocytes (Fig. 1b; P<0.00001), values again consistent with other studies (Chiang et al., 2010; Lister et al., 2010; Nakagawa and FitzHarris, 2017; Yun et al., 2014). Maternal aging thus increases the rate of anaphase lagging and aneuploidy, but the rate of lagging far exceeds that of aneuploidy. Of note, in addition to lagging chromosomes, our live imaging datasets also displayed previously reported chromosome segregation errors associated with cohesion loss, including balanced and non-balanced predivison (Sakakibara et al., 2015) and reverse segregation (Zielinska et al., 2015) (Fig. S2; *discussed further below*).

**Figure 1.**
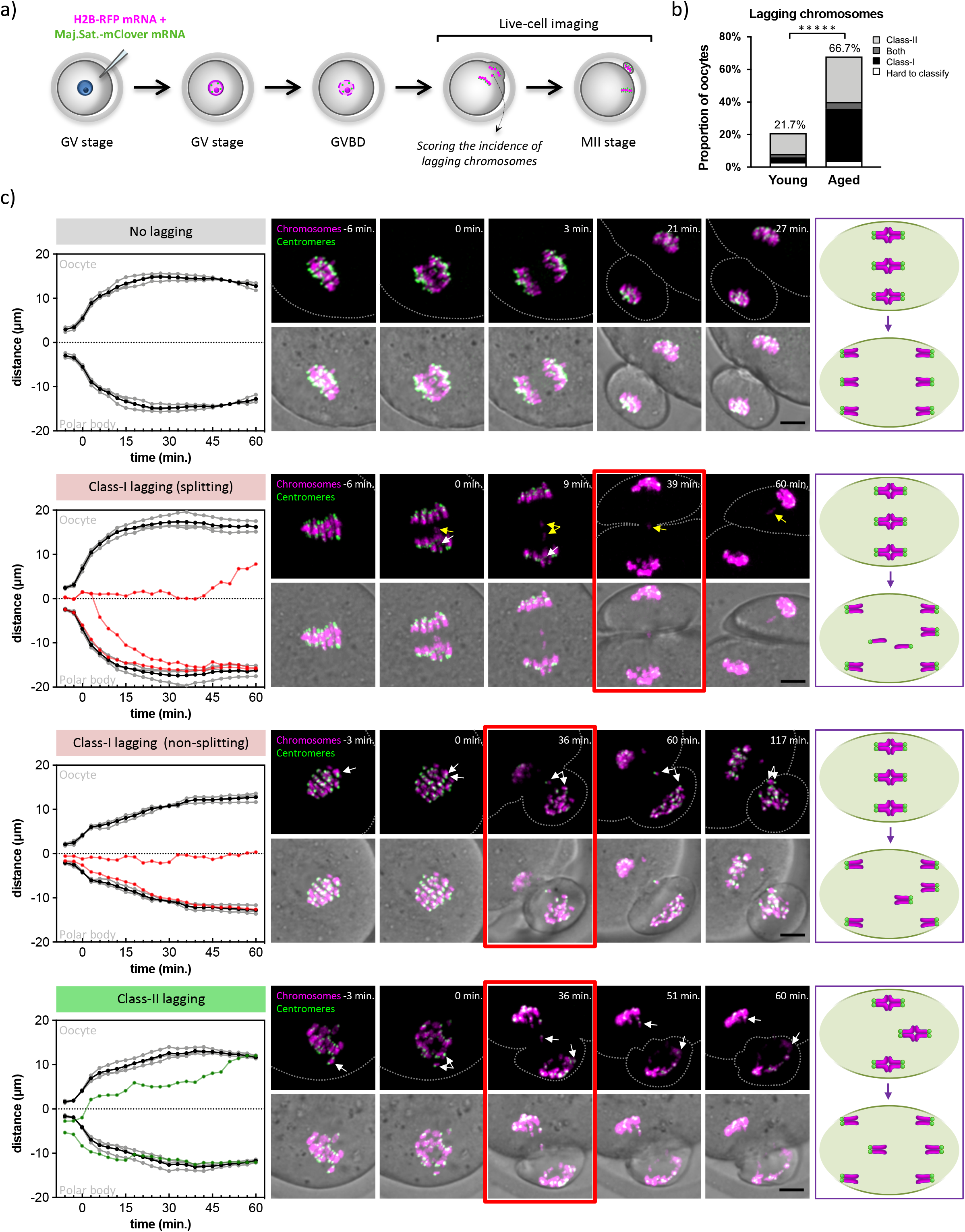
Two classes of lagging chromosomes. **a)** Experimental strategy to monitor lagging chromosome behavior during MI; **b)** The incidence of each lagging chromosome class in young (n=60) and aged (n=57) MI-stage oocytes (χ^2^-test, *****p<0.00001); **c)** Graphical representation of chromosome movement with corresponding time-lapse images showing anaphase with no lagging, Class-I and Class-II lagging chromosomes. Chromosomes are visualised with H2B-RFP (magenta) and centromeres with Maj.Sat.-mClover (green). Arrows indicate pairs of lagging sister chromatids and their non-lagging counterparts (white) or individual sister chromatids (yellow). Scale bars=20μm.

Next, we used our live imaging datasets to quantitatively describe the spatiotemporal dynamics of lagging chromosomes. Using 4D analysis of centromere position to monitor their individual trajectories, we discerned two distinct types of anaphase lagging chromosome behavior, that we refer to hereafter as ‘Class-I’ and ‘Class-II’. Class-I lagging chromosomes originated from fully congressed bivalents and exhibited reduced velocity of poleward movement during anaphase, causing the chromosome to lag behind in mid-anaphase (Fig. 1c), similar to how lagging is typically described in somatic cells. Contrastingly, a second class of laggard, which we term ‘Class-II’, originated from chromosomes that were very mildly misaligned at the time of anaphase onset (metaphase plate displacement 6.0 ± 0.2 μm) and moved apart in anaphase at a similar velocity to normally segregating chromosomes. However, the slight misalignment caused the sister pair facing the metaphase plate to lag behind other chromosomes in anaphase (Fig. 1c). Importantly, in the absence of temporal information the two types of lagging chromosomes would be completely indistinguishable from one another at mid-anaphase, both being evident as a chromosome in-between the two main masses of normally segregating chromosomes (Fig. 1c; see 36-39 min. timepoints in Class-I and Class-II examples for comparison). The incidence of both classes of laggards increased substantially with age (Fig. 1b; Class-I lagging 5.0% in young vs 35.1% in aged oocytes, P<0.00005; Class-II lagging 15.0% in young vs 31.6% in aged, P<0.04).

Having established the existence of two distinct classes of lagging made us investigate if these may have different outcome during MI. In a subset of movies we could continuously track the laggard and its non-lagging counterpart through the entirety of anaphase to their final destination. Strikingly, class-II laggard and its non-lagging counterpart were correctly shared between the oocyte and polar body in all cases (n=40). In sharp contrast, Class-I laggards were missegregated in 9 of 16 observable cases (~56%). Of those, the lagging pair of sister chromatids split during anaphase in 7 cases causing a gain or loss of an individual chromatid, while in the remaining 2 the intact pair missegregated into the wrong cell, indicating classical non-disjunction. These results thus reveal the existence of two distinct types of lagging chromosomes in mouse oocytes, wherein Class-I usually cause aneuploidy, but Class-II do not.

### M-phase prolongation reduces aneuploidy by eliminating class-I laggards

To further explore the relationship between lagging chromosomes and aneuploidy, we sought strategies to manipulate the occurrence of lagging. First, we extended M-phase using a reversible small molecule inhibitor of anaphase-promoting complex/cyclosome (APC/C), APCin (Sackton et al., 2014). We found that 100μM APCin potently inhibited the completion of MI, and its subsequent removal allowed resumption of MI and polar body extrusion within 18.5 ± 1.2 min. Having confirmed its reversibility, oocytes were exposed to a pulse of APCin to prolong MI by ~2.5h, from ~8.5h to ~11h in length. Chromosome behaviour was monitored to classify lagging chromosomes and *in situ* chromosome counting was subsequently performed on an oocyte-oocyte basis, allowing us to relate the class of lagging with aneuploidy outcome in the same oocyte (Fig. 2a). Overall, M-phase extension in aged oocytes significantly reduced both the incidence of lagging chromosomes (Fig. 2b, 63.3% in control vs. 25.7% in APCin group, P<0.01), and aneuploidy in resulting eggs (Fig. 2c, 36.7% in control vs. 14.3% in APCin group, P<0.04). Strikingly, the reduction of lagging was principally an eradication of Class-I laggards. In controls, 33.3% of oocytes exhibited Class-I lagging chromosome and almost all (9 out of 10) became aneuploid. In contrast, following M-phase extension only 2 oocytes (5.7%) had Class-I laggards, but both of these became aneuploid eggs. Class-II laggards were still a frequent occurrence in control and M-phase-extended oocytes, but oocytes with only class-II laggards very rarely (13.3%) became aneuploid. Thus, extending M-phase selectively suppressed the Class-I lagging chromosomes, and as a result reduced oocyte aneuploidy.

**Figure 2.**
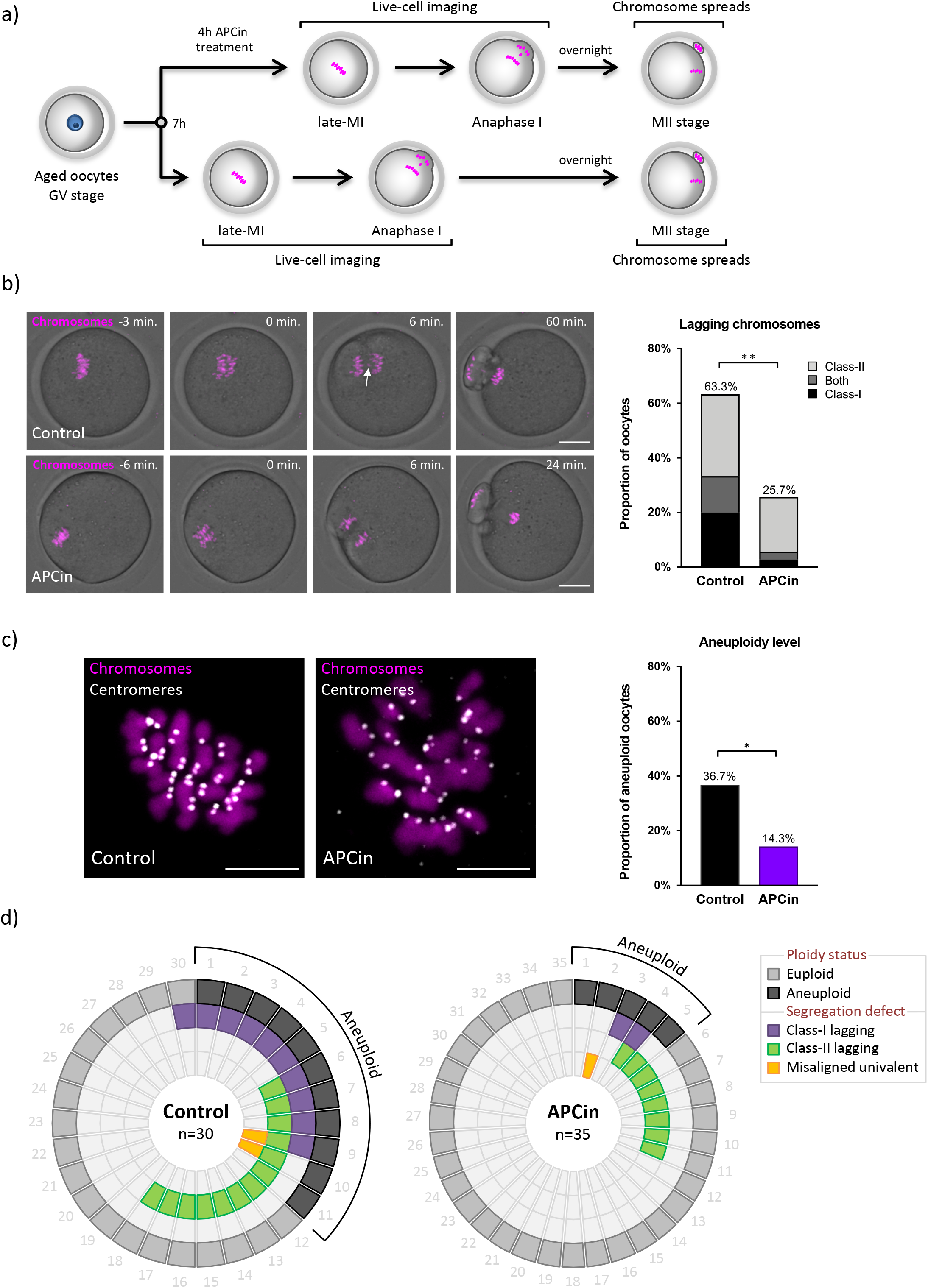
M-phase prolongation reduces Class-I lagging chromosome formation and aneuploidy in aged oocytes. **a)** Experimental strategy to assess lagging and ploidy status in individual oocytes after M-phase extension; **b)** Time-lapse confocal images of control (n=30) and APCin-treated (n=35) oocytes in anaphase. Chromosomes are labelled with SiR-DNA (magenta). White arrow denotes Class-l lagging chromosome. Note that the incidence of each lagging chromosome class in control was similar to aged group in Fig. 1b. Scale bars=20μm. The chart is showing the frequency of each lagging chromosomes class (χ^2^-test, **p<0.01). **c)** Exemplar *in situ* chromosome spreads of control (n=30) and APCin-treated (n=35) aged eggs with the chart showing aneuploidy level (χ^2^-test, *p<0.04). Chromosomes are labelled with Hoechst33342 (magenta) and centromeres with CENP-A antibody (grey). Scale bars=5μm; **d)** Doughnut diagrams show the correlation between chromosome segregation defects and aneuploidy in control and APCin-treated aged oocytes. The numbers around the diagram designate individual oocytes and ‘n’ is the total number of aged oocytes per group.

Correct microtubule (MT) attachment to kinetochores is central to accurate chromosome segregation and avoidance of chromosome lagging in somatic cells. Therefore, to understand why laggards are common in aged oocytes, and how M-phase prolongation reduced Class-I lagging chromosomes, we explored how kinetochore-microtubule (KT-MT) attachments are established in MI in young and aged oocytes, using cold-shock treatment to visualise stable MTs (Fig. 3a). In both young and aged oocytes, the proportion of end-on KT-MT attachments gradually increased over the course of MI (from ~10% to ~65-95%), and the proportion of lateral ‘side-on’ KT-MT attachments decreased correspondingly (from initial ~60-70% to ~5-15%; fig. 3b). In both groups, the proportion of merotelic attachments was highest 3h post-MI entry (6.0% and 8.4% in young and aged, respectively), but then gradually decreased and reached negligible levels shortly prior anaphase (Fig. 3b; 0.6% vs. 1.3% in young vs. aged group at 8h post-MI entry, respectively). Thus, merotelic attachments are no more abundant in aged oocytes, and their resolution occurs with apparently normal kinetics. Rather, the major difference in KT-MT attachment status between two groups was that aged oocytes possess a major pool of ~30% of all kinetochores that remain entirely unattached even late in MI. In contrast, almost no kinetochores remained unattached at 8h post-MI entry in young oocytes (0.3%). These data suggest that aging reduces chromosomes capacity to establish stable k-fibres, resulting in a significant pool of unattached kinetochores in late MI oocytes from aged animals.

**Figure 3.**
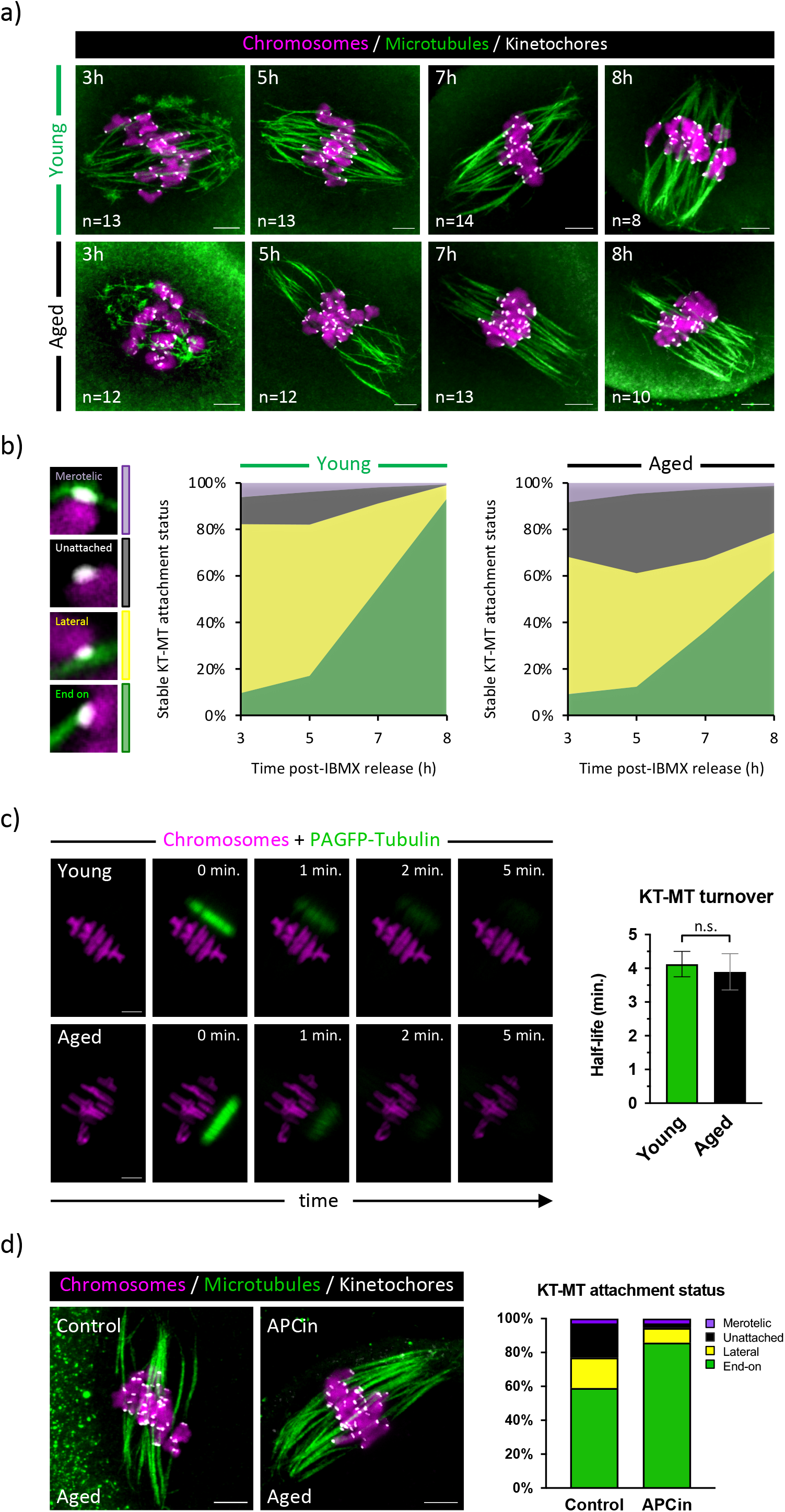
The age-related changes in k-fibre formation during MI. **a)** Exemplar confocal images of coldshock treated young and aged oocytes 3, 5, 7 and 8h post-IBMX release. Chromosomes are labelled with Hoechst33342 (magenta), microtubules with β-tubulin antibody (green) and kinetochores with CREST (grey); ‘n’ is the total number of oocytes per group. Scale bars=5μm; **b)** The proportion of different KT-MT attachment statuses (end-on, lateral, unattached and merotelic) in young and aged MI-oocytes at designated time-points; **c)** Exemplar confocal images of young (n=23) and aged (n=18) late-MI oocytes showing dissipation of PAGFP-Tubulin fluorescent signal. Scale bars=5μm. The graph shows KT-MT turnover rate comparison between young and aged oocytes (unpaired t-test, p=0.22); **d)** Exemplar confocal images of cold-shock treated control (n=7) and APCin-treated (n=9) aged oocytes. Chromosomes are labelled with Hoechst33342 (magenta), microtubules with β-tubulin antibody (green) and kinetochores with CREST (grey). Scale bars=5μm. The chart shows proportion of KT-MT attachment statuses in each group.

To test this conclusion with a distinct experimental approach, we employed fluorescence dissipation after photoactivation (FDAP) of photoactivatable-GFP:Tubulin (PAGFP-Tubulin). We created a line of photoactivated signal between the chromosomes and one spindle pole in young and aged oocytes and live-imaged its dissipation rate (Fig. 3c) to construct FDAP decay curve comprised of two components. A slow turnover component (T1/2 in min.) provides a readout of KT-MT turnover that is necessary for error correction, whereas a fast turnover component (T1/2 in sec.) indicates non-stabilised spindle MTs (Bakhoum et al., 2009a; Cimini et al., 2006; Paim and FitzHarris, 2019; Zhai et al., 1995). Mathematical deconvolution of these two components reveals the relative sizes of the two MT populations (Fig. S3). Strikingly, we observed no significant difference in slow turnover component between young and aged oocytes (Fig. 3c and S3; T1/2 = 4.1 ± 0.4 min. in young vs. T1/2 = 3.9 ± 0.6 min. in aged, p=0.22), consistent with the similar error-correction rates measured in Fig. 3b. In line with our previous finding, the proportion of slow turnover component which corresponds to stable k-fibres was lower in aged oocytes (Fig. S3; 16.8% in aged vs 22.1% in young). Thus, both approaches support the conclusion that there is not an appreciable age-related deficit in MT error-correction mechanisms but rather k-fibre establishment is slowed in aged oocytes.

We wondered how M-phase extension reduced lagging and aneuploidy, and reasoned that M-phase extension likely provided more time for stable k-fibre establishment. To test this, we assessed KT-MT attachments in M-phase prolonged oocytes. Strikingly, the proportion of chromosomes with unattached kinetochores was reduced to a negligible level (Fig. 3d; 2.6% in M-phase-prolonged vs 20.1% in control) while the proportion of merotelic attachments remained almost identical as in untreated oocytes (Fig. 3d; 2.8% vs. 2.9%). Taken together, this series of experiments shows that establishment of k-fibres is slowed in aged oocytes, but M-phase extension provides enough extra time to establish correct KT-MT attachments, thereby lowering the incidence of Class-I lagging chromosomes that cause oocyte aneuploidy.

### MCAK replacement reduces class-II lagging without impacting aneuploidy

Next, we sought an entirely different approach to decrease the incidence of lagging chromosomes. In mitotic cells, depletion of mitotic centromere-associated kinesin (MCAK) increases production of lagging chromosomes (Ganem et al., 2005; Kline-Smith et al., 2004; Maney et al., 1998) while its overexpression can reduce the likelihood of lagging (Bakhoum et al., 2009b; Kabeche and Compton, 2012; Paim and FitzHarris, 2019; Wordeman et al., 2007). In oocytes, MCAK transcript levels decrease with maternal age (Pan et al., 2008), causing us to hypothesize that it may underlie age-related anaphase lagging. To explore this possibility, we first examined whether MCAK protein is dysregulated with age and observed a clear age-related decrease in centromeric/kinetochore MCAK (Fig. 4a; 42% reduction, P<0.0001). To determine whether supplementing aged oocytes with MCAK could reduce lagging, we overexpressed MCAK-GFP or GFP and observed chromosomes behaviour during MI (Fig. 4b). MCAK-GFP overexpression significantly reduced the occurrence of lagging chromosomes (Fig. 4c, 62.1% vs. 36.8% in GFP vs. MCAK-GFP group, P<0.05). Notably, analysis of laggard types showed that the frequency of Class-I laggards remained unchanged, but Class-II laggards were lessened (Fig. 4c; 41% in GFP vs. 11% in MCAK-GFP group, P<0.004). Importantly, the aneuploidy level remained unchanged after MCAK-GFP overexpression (Fig. 4d; 30.4% in control vs. 28.0% in MCAK-GFP aged oocytes, P>0.85). Thus, whereas M-phase extension limited Class-I lagging chromosome number and reduced aneuploidy, MCAK overexpression decreased Class-II lagging chromosomes and had no impact on oocyte aneuploidy.

**Figure 4.**
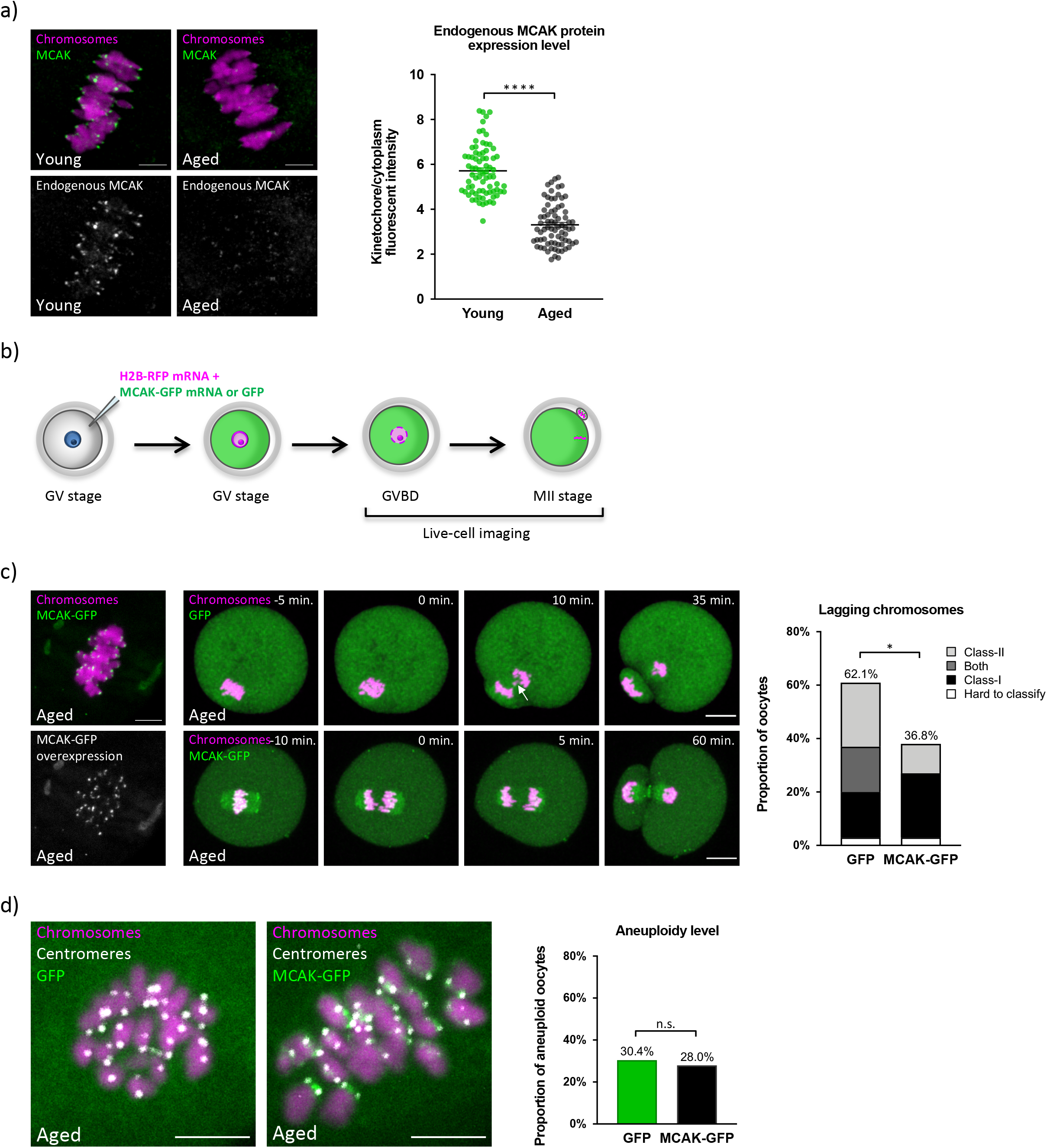
The age-associated MCAK reduction contributes towards excessive Class-II lagging chromosome formation but not aneuploidy. **a)** Exemplar confocal images of young (n=15) and aged (n=15) late-MI oocytes, showing the endogenous MCAK protein expression level (green) and chromosomes (magenta), labelled with MCAK antibody and Hoechst33342, respectively (top panels). The endogenous MCAK protein expression is shown separately in grey scale (bottom panels). Scale bars=5μm. Quantification of the endogenous MCAK protein (unpaired t-test, ****p<0.0001); **b)** Experimental strategy to assess the incidence of lagging chromosomes in MCAK-GFP and GFP overexpressing aged oocytes during MI; **c)** Overexpressed MCAK-GFP protein localizes at the kinetochore (n=21); MCAK-GFP native fluorescent signal is shown combined with (green) or separately from (grey) Hoechst33342-labelled chromosomes (magenta). Scale bars=5μm. Middle panels show time-lapse confocal images of MCAK-GFP or GFP overexpressing (green) aged oocytes during anaphase. Chromosomes are visualised with H2B-RFP (magenta). White arrow denotes Class-II laggard. Scale bars=20μm. The proportion of each lagging chromosome class in MCAK-GFP (n=52) and GFP (n=47) overexpressing aged oocytes (χ^2^-test, *p<0.05); **d)** Exemplar *in situ* chromosome spreads of MCAK-GFP (n=25) and GFP (n=23) aged eggs with the chart showing aneuploidy level (χ^2^-test, p>0.85). Chromosomes are labelled with Hoechst33342 (magenta), centromeres with CENP-A antibody (grey) and merged with MCAK-GFP or GFP native fluorescence signal (green). Scale bars=5μm.

## DISCUSSION

In the present study we show that not all laggards are equal. Instead, we identify two mechanistically distinct types of lagging chromosomes in MI that, though appearing identical in mid-anaphase, have different outcomes on chromosome segregation. Whilst the type we have dubbed Class-II are almost never ultimately missegregated, Class-I are often missegregated and thus cause aneuploidy. Our data thus shows that lagging chromosomes provide a direct route to aneuploidy, and simultaneously resolves the paradox as to why lagging chromosomes are almost always in excess of aneuploidy in oocytes.

Our findings complement the well-established notion that chromosome cohesion decline is a major contributor to age-related aneuploidy (Sakakibara et al., 2015). Although we focus here on lagging chromosomes, our imaging datasets also reveal all of the previously reported cohesion loss associated defects (Chiang et al., 2010; Sakakibara et al., 2015; Zielinska et al., 2015). Specifically, we observed premature separation of bivalents into univalents in 19.3% of aged oocytes (Fig. S2a), which either led to gains or losses of single chromatids in ~2/3 of cases (i.e. non-balanced predivision) or balanced predivision (Fig. S2b). The occurrence of misaligned univalent shortly prior anaphase always led to aneuploidy as it traveled intact to the spindle pole while its aligned counterpart segregated sister chromatids causing nonbalanced predivision (Fig. S2c). In addition, for the first time, we report in mouse an example of reverse segregation (Fig. S2d), previously observed only in human (Zielinska et al., 2015), wherein partial separation of sister kinetochores allows the bivalent rotated by 90 degrees to align at the metaphase plate. We also observed true non-disjunction (Fig. S2e) when severely misaligned bivalent segregated intact towards the nearest spindle pole (Sodek et al., 2017). Our data thus confirm previous reports that cohesion loss predisposes chromosomes to missegregation in aged oocytes and formally add the lagging chromosome (specifically Class-I) as a previously undefined route to aneuploidy.

Could lagging chromosomes be a knock-on effect of cohesion loss? Whilst age-related cohesion loss has been argued to favour the formation of merotelic attachments (Chiang et al., 2010; Zielinska et al., 2019), the observed correlation between KT-MT attachments status and cohesion loss in mouse oocytes is weak (Shomper et al., 2014). Our finding that M-phase extension or MCAK overexpression, manipulations with no obvious effect on chromosome cohesion, can reduce lagging chromosome formation additionally argues against this possibility (Fig. 2b and 4c). Moreover, lagging chromosomes are seen in young oocytes with full cohesion complement (Nakagawa and FitzHarris, 2017; Yun et al., 2014) and age-associated alterations in MT dynamics documented here (Fig. 3) and elsewhere (Nakagawa and FitzHarris, 2017) provide a mechanistic explanation for their increase in aged oocytes. Overall, our data indicate that chromosome lagging is an age-related lesion that directly causes aneuploidy in mammalian oocytes independently of chromosome cohesion loss.

One apparent reason for the increase in lagging chromosome formation in aged oocytes is an age-associated reduction in centromere/kinetochore MCAK expression. However, MCAK appears to prevent Class-II lagging chromosomes that do not generally cause aneuploidy. Although this may seem unexpected given the role of MCAK in KT-MT error-correction in mitotic cells (Kline-Smith et al., 2004; Knowlton et al., 2006; Sanhaji et al., 2011; Wordeman et al., 2007), it is in direct agreement with previous reports that MCAK dysregulation in oocytes causes chromosome misalignment but not segregation error (Illingworth et al., 2010; Vogt et al., 2010). It is also consistent with the relative paucity of merotelic attachments found in young or aged oocytes (Fig. 3b). Thus MCAK-mediated error correction, central to aneuploidy avoidance in somatic cells, is largely dispensable in oocytes wherein bivalents are positioned prior to the establishment of k-fibres, and true merotelic attachments are extremely rare.

Finally, our study shows that extension of M-phase can reduce the incidence of lagging chromosomes and age-related aneuploidy in mouse oocytes. Whilst extensive tests of efficacy and safety are necessary before such an approach was ever to be used clinically, our data raise the intriguing possibility that a simple chemical intervention, such as modulation of M-phase length, could eventually be used in clinical context to improve the egg quality of patients at advanced age.

## SUPPLEMENTARY FIGURE LEGENDS

**Supplementary Figure 1.**
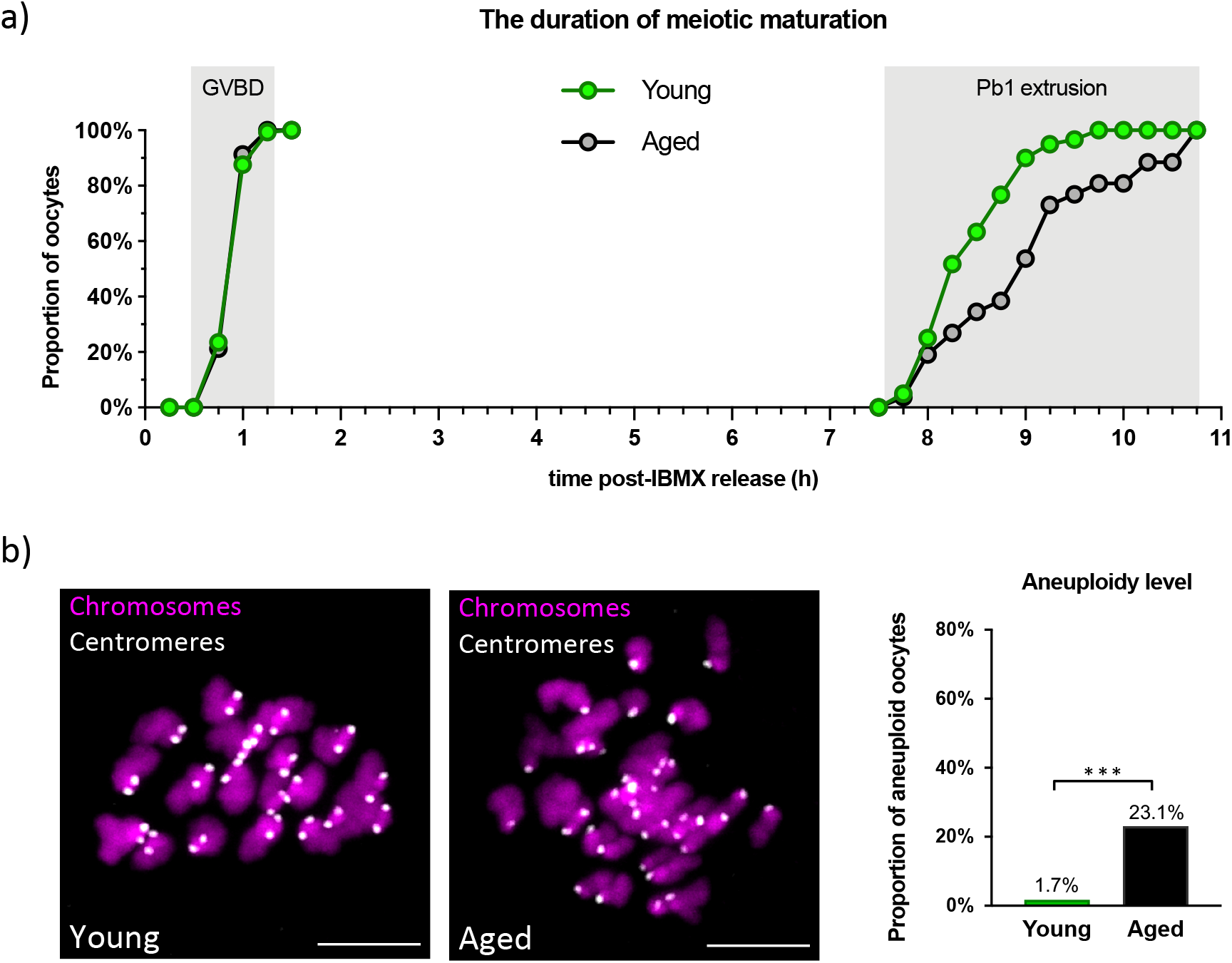
The age-associated increase in aneuploidy. **a)** *In vitro* meiotic maturation progression in young (n=60) and aged (n=26) oocytes detailing the rate of germinal vesicle breakdown and the first polar body extrusion; **b)** Exemplar *in situ* chromosome spreads of young (n=60) and aged (n=26) eggs with the chart showing aneuploidy level (χ^2^-test, ***p<0.001). Chromosomes are labelled with Hoechst33342 (magenta) and centromeres with CENP-A antibody (grey). Scale bars=5μm.

**Supplementary Figure 2.**
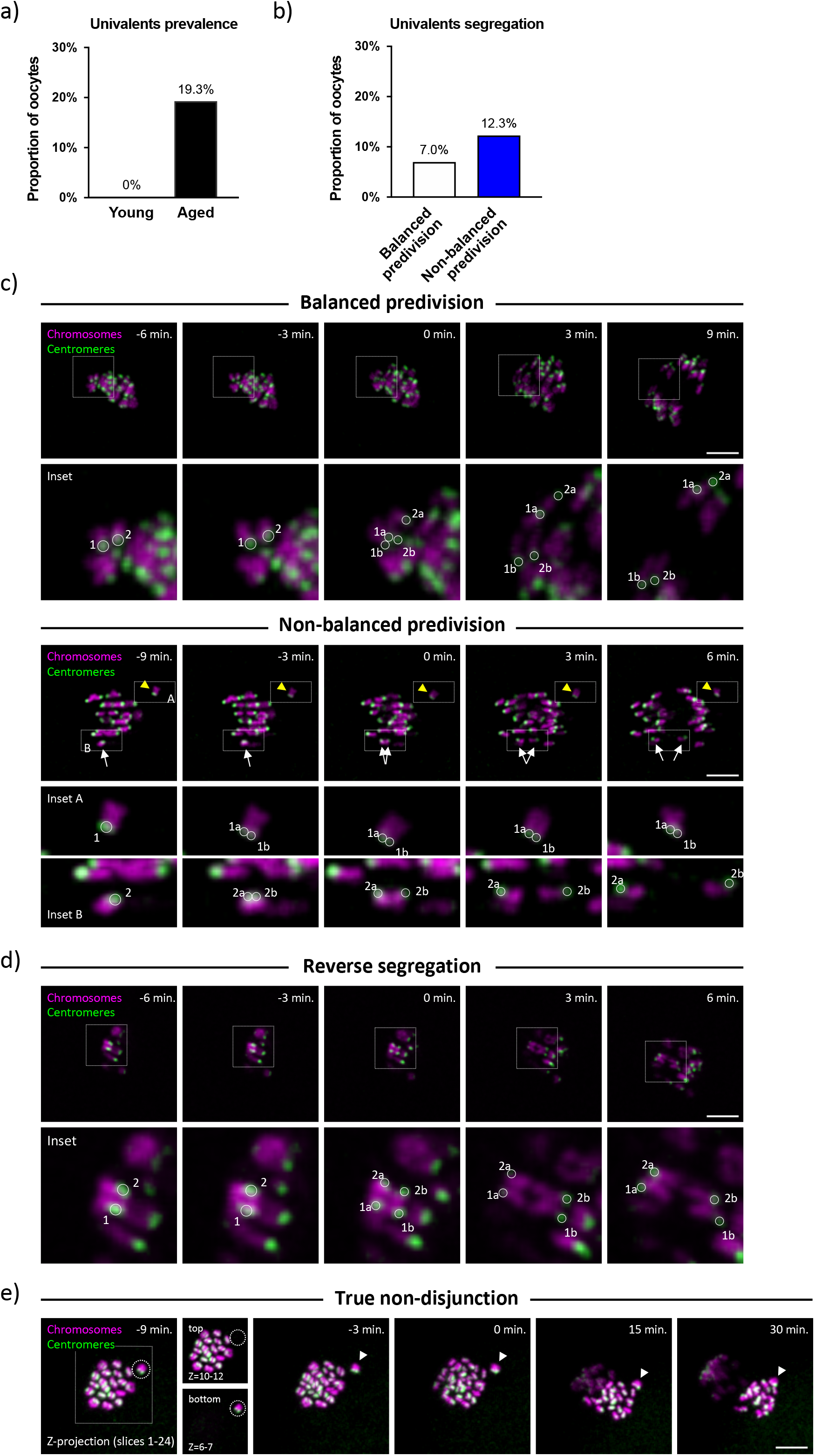
Previously reported chromosome segregation errors. **a)** The incidence of univalents formation in young (n=60) and aged (n=57) MI-oocytes; **b)** The univalents segregation outcome in aged oocytes; **c)** Examples of balanced (top) and non-balanced (bottom) predivsion in aged oocytes. Insets detail univalents segregation; numbers (1,2) and letters (a,b) denote univalents and segregating sister chromatids, respectively. Note that sister chromatids of aligned univalents (white arrows) separate towards opposite spindle poles while misaligned univalent (yellow arrowhead) travels intact towards the nearest spindle poles causing aneuploidy. **d)** Reverse segregation - sister chromatids of 90° rotated bivalent split evenly between the oocyte and the polar body. Inset details separation of sister chromatids, numbers (1,2) and letters (a,b) denote pair of sister chromatids and individual chromatids, respectively. **e)** A severely misaligned bivalent travels intact towards the nearest spindle pole causing true non-disjunction. Insets are partial Z-projections detailing the plane of the metaphase plate (top) and misaligned bivalent (bottom). Dashed circle line marks the position of misaligned bivalent in Z-axis and the arrowhead denotes misaligned bivalent at subsequent time-points. In **c)**, **d)** and **e)**, chromosomes are labelled with H2B-RFP (magenta) and centromeres Maj.Sat.-mClover (green). The indicated time-points are relative to the anaphase onset. Scale bars=10μm.

**Supplementary Figure 3.**
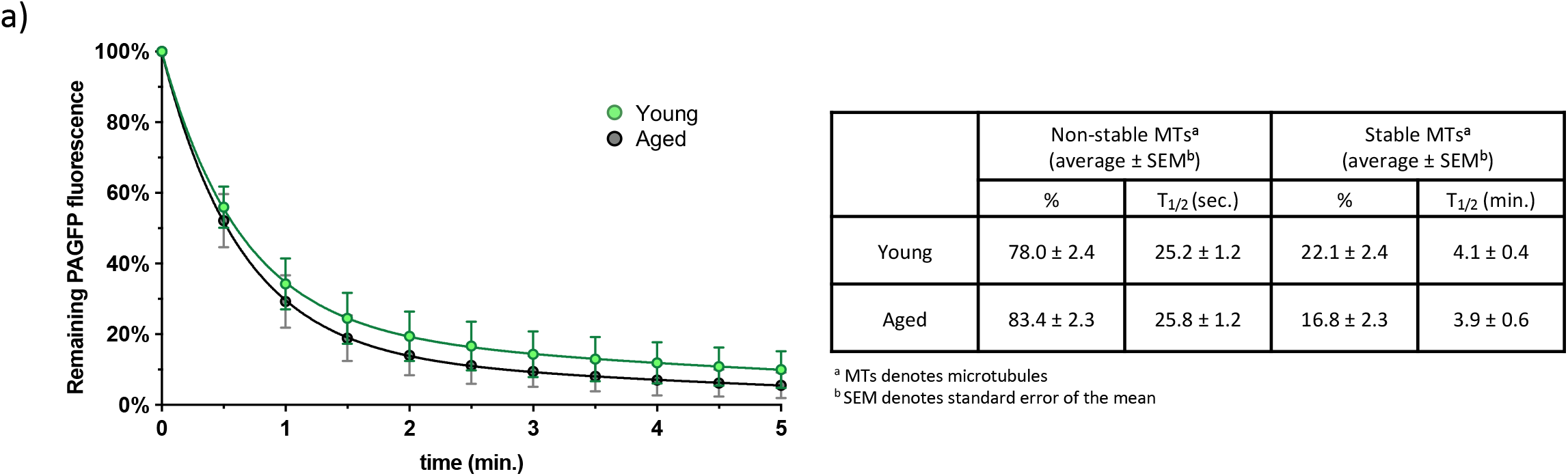
Quantification of PAGFP-Tubulin fluorescence decay rate in late-MI oocytes. The fluorescence dissipation after photoactivation curve represents the average fluorescence decay rate of PAGFP-Tubulin in young (n=23) and aged (n=18) oocytes late in MI. The presented values are mean ± SEM per each time-point.

## EXPERIMENTAL PROCEDURES

### Oocyte collection and handling

All experiments were approved by the Centre Hospitalier de l’Université de Montréal (CRCHUM) Comité Institutionnel de Protection des Animaux du CHUM (CIPA). Fully grown, germinal vesicle (GV)-stage oocytes were collected from 2-3 and 16 months of age CD1 mice, 44-46h following pregnant mare’s serum gonadotrophin (PMSG) hormonal stimulation (5 and 10 IU of PMSG/mouse, respectively). Oocyte handling outside the incubator was done in M2 media and in the presence of 200uM 3-isobutyl-1-methylxanthine (IBMX) to maintain them at GV stage. The surrounding cumulus cells have been removed and the oocytes washed through M16 media with 200μM IBMX and left in the incubator at 37°C and 5% CO2 for approximately 1h to recover prior any further manipulation.

### Microinjection

GV-stage oocytes were microinjected in M2 media supplemented with with 200μM IBMX, using a picopump (World Precision Instruments), intracellular electrometer (Warner instruments), and micromanipulators (Narishige), mounted on a Leica DMI4000 inverted microscope (as described previously Fitzharris, 2009; FitzHarris et al., 2018). The mRNAs used for injections were *in vitro* synthesised using mMessage mMachine kits (T3, T7 and SP6), followed by poly-adenylation using Poly(A)-Tailing kit according to the manufacturer’s instructions. The following linearised plasmids were used as templates: pRN3:H2B-RFP, pTALYM3B15 (Maj.Sat./TALE-mClover), pIRESHyg2:human-α-Tubulin-paGFP, pCS2mt:GFP and pcDNA3.1/*myc*-His(-)A:MCAK-GFP. Injected oocytes were left in the incubator for at least 3 hours to allow the time for fluorescent proteins expression.

### Live-cell confocal imaging

GV-stage oocytes were released from the IBMX inhibition into M16 media to allow the resumption of meiotic maturation and placed in an on-stage incubating chamber (at 37°C and 5% CO2) that was mounted on a Leica SP8 confocal microscope. Z-stack were acquired with 20x air objective (0.75 NA) using 2μm optical section and step size at 3 min time-interval for live-imaging experiments in Fig. 1c and 2b, and 5min time-interval in Fig. 4c. For the experiments in Fig. 1c and 4c, the oocytes were illuminated with 488nm and 552nm light (using 0.5% and 0.15% power, respectively) and in Fig. 2b with 638nm laser (0.25% power) and fluorescence was detected with a HyD detector. For Fig. 3c, a single Z-section time-lapse images were obtained at 30 second time-intervals.

### APCin treatment

To extend the M-phase, aged GV-stage oocytes were first allowed to resume meiotic maturation in (IBMX-free) M16 media, and at 7h following their release from IBMX were transferred into M16 with 100μM APCIN for 4h. APCin was then either washed out and oocytes live-imaged, or oocytes were immediately exposed to cold-shock prior to fixation. For live-imaging experiments in Fig. 2, oocytes were incubated in M16 with 1μM SiR-DNA for 2h to label the chromosomes.

### Fixation and immunofluorescence

Oocytes were fixed using 2% PFA in PBS for 20 min, followed by permeabilization step in 0.25% Triton X-100 in PBS for 10 min at ambient temperature. Blocking was perfomed in 3% BSA for 1h or overnight at 4°C. For experiments where *in situ* chromosome counting was performed (Fig. 2c, 4d and Fig. S1), the oocytes were pre-treated with λ-protein phosphatase for 30 min at 30°C according to manufacturer’s instructions. The following primary antibodies were used in this study: rabbit anti-CENP-A (1:200), mouse anti-β-tubulin (1:1000), rabbit anti-MCAK (1:1000) and CREST (1:100). Secondary antibodies as follows: goat anti-rabbit Alexa488 (1:1000), goat anti-rabbit Alexa568 (1:1000), donkey anti-rabbit Alexa647 (1:1000), goat anti-mouse Alexa488 (1:1000) and goat anti-human Alexa546 (1:1000). Chromosomes were labelled with Hoechst 33342 (1:500) for 20 min at ambient temperature. Imaging of fixed samples was carried out on a Leica SP8 confocal microscope using 63xoil-immersion objective (1.4 NA) using 0.9μm optical section and step size. To assess cold-stable KT-MT attachments, culture dishes containing oocytes were exposed to wet ice for 10 min immediately prior to fixation. For the experiments were ploidy status was assessed by *in situ* chromosome counting (Fig. 2c, 4d, S1), following completion of meiotic maturation the oocytes were treated with 200μM Monastrol in M16 media for 2h prior fixation.

### Microtubule turnover measurement

To measure the microtubule turnover rates in Fig. 3c and Fig. S3, fluorescence dissipation after photoactivation (FDAP) was performed by photoactivating paGFP-Tubulin with 405nm laser within a defined rectangular shaped region of interest positioned in between the metaphase plate and one of the spindle poles and live imaging subsequently performed at 30 sec. time intervals for 5 minutes. The fluorescence decay curves and MT half-live (T1/2) were obtained by plotting the fluorescence intensity decay obtained for each spindle against time and fitted into a double exponential curve *f(t)=A x exp(-k_1_t) + B x exp(-k_2_t)* using the cftool in MATLAB, wherein *t* represents the time; A is the non-stable MT population, B is the stable MT population (KT-MT) and k1 and k2 are the decay rates of the two MT populations, respectively. The half-life (T1/2) of KT-MT population was calculated as ln2/k2. Each measurement was corrected for photobleaching. The extent of photobleaching was measured by imaging young late-MI stage oocytes exposed to 10μM MT-stabilising agent, Taxol, where the MT-turnover rate was minimal.

## QUANTIFICATION AND STATISTICAL ANALYSIS

### Image analysis and statistics

Image analysis was performed in ImageJ/Fiji and Imaris (version 9.6.0, Bitplane). Anaphase chromosome was considered lagging if it was optically separate from the main body of segregating chromosomes when the spindle was oriented to be planar to the field of view. For Fig. 1c, spatial coordinates of lagging and referent chromosomes (3 normally segregating bivalents per example) during anaphase were manually determined based on Maj.Sat.-mClover signal and the distance between segregating chromosomes was calculated using Pythagoras’ theorem. For Fig. 4a, MCAK immunofluorescence signal intensity was quantified by calculating the ratio between the average fluorescent signal (mean grey value) of individual kinetochores and cytoplasmic background from a total of 75 kinetochores per group (15 young and 15 aged; 5 kinetochores per oocyte analysed). Statistical analysis was performed using GraphPad Prism8 software using χ^2^-test and unpaired t-test where appropriate. Error bars represent as standard error of the mean.

## KEY RESOURCES TABLE

| REAGENT or RESOURCE | SOURCE | IDENTIFIER |
| --- | --- | --- |
| Antibodies |  |  |
| Rabbit anti-CENP-A | New England Biolabs | Cat# 2048S; RRID: AB_1147629 |
| Mouse anti-β-tubulin | Sigma | Cat# T4026; RRID: AB_477577 |
| CREST | A kind gift from Marvin J. Fritzler | N/A |
| Rabbit anti-MCAK | (Mack and Compton, 2001) | N/A |
| Alexa Fluor 488 anti-Mouse | ThermoFisher | Cat# A-11029; RRID: AB_2534088 |
| Alexa Fluor 488 anti-Rabbit | ThermoFisher | Cat# A-11008; RRID: AB_2540618 |
| Alexa Fluor 546 anti-Human | ThermoFisher | Cat# A-21089; RRID: AB_2535745 |
| Alexa Fluor 647 anti-Rabbit | ThermoFisher | Cat# A-31573; RRID: AB_2536183 |
| Alexa Fluor 568 anti-Rabbit | ThermoFisher | Cat# A-11011; RRID: AB_143157 |
| Chemicals, Peptides and Recombinant Proteins |  |  |
| Pregnant Mare's Serum Gonadotropin (PMSG) | Genway Biotech | Cat# GWB-2AE30A |
| M2 Media | Sigma | Cat# M7167 |
| M16 Media | Sigma | Cat# M7292 |
| IBMX (3-Isobutyl-1-methylxanthine) | Sigma | Cat# I5879; CAS: 28822-58-4 |
| APCin | Tocris | Cat# 5747; CAS: 300815-04-7 |
| Monastrol | Sigma | Cat# 475879; CAS: 254753-54-3 |
| Taxol | Sigma | Cat# T7402; CAS: 33069-62-4 |
| SiR-DNA | Cytoskeleton Inc. | Cat# CY-SC007 |
| Hoechst33342 | Invitrogen | H1399 |
| λ-protein phosphatase | New England Biolabs | Cat# P0753S |
| Paraformaldehyde | Sigma | Cat# P6148; CAS: 30525-89-4 |
| Triton X-100 | Sigma | Cat# T9284; CAS: 9002-93-1 |
| Bovine Serum Albumin (BSA) | Sigma | Cat# A7906; CAS: 9048-46-8 |
| **Critical Commercial Assays** |  |  |
| mMessagemMachine Kit (SP6, T3, T7) | Ambion | Cat# AM1340, AM1348, AM1344 |
| Poly(A) Tailing Kit | Ambion | Cat# AM1350 |
| **Experimental Models** |  |  |
| CD-1 IGS mice | Charles River Laboratories | CrI:CD1 (ICR) |
| ICR (CD-1) outbred mice | Envigo | Hsd:ICR (CD-1) |
| **Recombinant DNA** |  |  |
| pRN3:H2B-RFP | (Prodon et al., 2010) | N/A |
| pTALYM3B15 (Maj.Sat.-mClover) | (Miyanari et al., 2013) | Addgene Plasmid #47878 |
| pIRESHyg2: human-α-Tubulin-paGFP | (Tulu et al., 2003) | Addgene Plasmid #12296 |
| pCS2mt: GFP | N/A | Addgene Plasmid #15681 |
| pcDNA3.1/myc-His(-)A: MCAK-GFP | (Illingworth et al., 2010) | N/A |
| **Software and Algorithms** |  |  |
| Fiji | (Schindelin et al., 2012) | https://imagej.net/Fiji |
| Imaris | Bitplane | https://imaris.oxinst.com |
| GraphPad Prism | GraphPad Prism8 Software, La Jolla California USA | https://www.graphpad.com |

## SUPPLEMENTAL INFORMATION

Supplemental information includes three figures and a table.

## AUTHOR CONTRIBUTIONS

A.I.M. and G.F. conceptualised the study. A.I.M. and J.H. performed the experiments and analysed the data. A.I.M. prepared the figures. A.I.M. and G.F. wrote the paper.

## ACKNOWLEDGMENTS

We thank Gaudeline Remillard-Labrosse for excellent laboratory support and Caitlin Mehrotra for assistance in data analysis. We also thank John Carroll and Gary Brouhard for critical reading of the manuscript. This research was supported by the grants to G.F. from Canadian Institute for Health Research (CIHR) and Foundation Jean-Louis Levesque. A.I.M. was supported by the Fonds de Recherche du Québec-Santé (FRQS) and McGill’s Centre for Research in Reproduction and Development (CRRD) Postdoctoral Fellowships.

